# Functional characterization of SARS-CoV-2 infection suggests a complex inflammatory response and metabolic alterations

**DOI:** 10.1101/2020.06.22.164384

**Authors:** Lucía Trilla-Fuertes, Ricardo Ramos, Natalia Blanca-López, Elena López-Camacho, Laura Martín-Pedraza, Pablo Ryan Murua, Mariana Díaz-Almirón, Carlos Llorens, Toni Gabaldón, Andrés Moya, Juan Ángel Fresno Vara, Angelo Gámez-Pozo

## Abstract

Covid-19, caused by the SARS-CoV-2 virus, has reached the category of a worldwide pandemic. Even though intensive efforts, no effective treatments or a vaccine are available. Molecular characterization of the transcriptional response in Covid-19 patients could be helpful to identify therapeutic targets. In this study, RNAseq data from peripheral blood mononuclear cell samples from Covid-19 patients and healthy controls was analyzed from a functional point of view using probabilistic graphical models. Two networks were built: one based on genes differentially expressed between healthy and infected individuals and another one based on the 2,000 most variable genes in terms of expression in order to make a functional characterization. In the network based on differentially expressed genes, two inflammatory response nodes with different tendencies were identified, one related to cytokines and chemokines, and another one related to bacterial infections. In addition, differences in metabolism, which were studied in depth using Flux Balance Analysis, were identified. SARS-CoV2-infection caused alterations in glutamate, methionine and cysteine, and tetrahydrobiopterin metabolism. In the network based on 2,000 most variable genes, also two inflammatory nodes with different tendencies between healthy individuals and patients were identified. Similar to the other network, one was related to cytokines and chemokines. However, the other one, lower in Covid-19 patients, was related to allergic processes and self-regulation of the immune response. Also, we identified a decrease in T cell node activity and an increase in cell division node activity. In the current absence of treatments for these patients, functional characterization of the transcriptional response to SARS-CoV-2 infection could be helpful to define targetable processes. Therefore, these results may be relevant to propose new treatments.

## Introduction

The emerging coronavirus SARS-CoV-2 has rapidly expanded from its origin in Wuhan, China, to become a worldwide pandemic only after four months since its first identification. At September 20^th^ of 2020, 30,675,675 cases and 954,417 deaths have been reported worldwide, according to the World Health Organization [1].

The most common symptoms are fever, cough, fatigue, shortness of breath, accompanied by elevated inflammatory biomarkers and pulmonary infiltrates. However, during the SARS-CoV-2 infection, a fraction of patients will develop severe pneumonia, pulmonary oedema, severe acute respiratory syndrome (SARS) or multiple organ failure, ending in death [2]. These severe symptoms are associated with systemic inflammation related to an overproduction of macrophagic cytokines. Different treatments focused on these inflammatory processes are being investigated [3].

Recently, Xiong et al. analyzed the transcriptional response in samples from peripheral blood mononuclear cells (PBMCs) from patients diagnosed with SARS-CoV-2 and compared them with healthy controls. Based on the results, the authors suggested that patient’s lymphopenia may be caused by an activation of apoptosis in lymphocytes, and also that SARS-Cov-2 induced excessive cytokine production, which correlates with lung tissue injury [4]. However, these conclusions were based only in the functional enrichment analysis of differentially expressed genes.

Probabilistic graphical models (PGMs) have demonstrated their utility in analyzing gene expression data by identifying relevant biological processes [5, 6]. These models allow making associations between genes according to their expression patterns across a series. Interestingly, the PGM networks have functional structure, allowing study expression data from a functional point of view. The main advantage of this type of models is that they offer an integrated view about what biological processes are involved in a disease, instead of the classical gene-based analysis which offers a list of differential genes without a context. Thus, we set out to re-analyze Xiong et al. data using PGMs, aiming for a deeper understanding of biological processes involved in SARS-CoV-2 pathogenesis.

## Results

### Processing of RNA sequencing data

After alignment of raw files, 13,398 expressed genes were identified. After applying the quality criteria of a detectable reading in at least 50% of the samples, 13,182 genes were used for the subsequent analyses.

### Analysis of differential genes between healthy controls and patients

Using CuffDiff, 1,569 differentially expressed genes were determined between SARS-CoV-2 patients and healthy controls. After applying the quality criteria of detectable measurements in at least 50% of the samples, 1,234 genes remained as differential ones. These genes were mostly related to inflammatory response, innate immune response, T cells, lysosomes, apoptotic processes and angiogenesis, among others.

A PGM was used to organize these genes according to their biological functions. The resulting network was composed by eight functional nodes: metabolism, lysosomes, T cells, two nodes related to inflammatory response, two nodes related to response to virus, and one node with no overrepresented function (Fig 1, S1 File).

**Fig 1:**
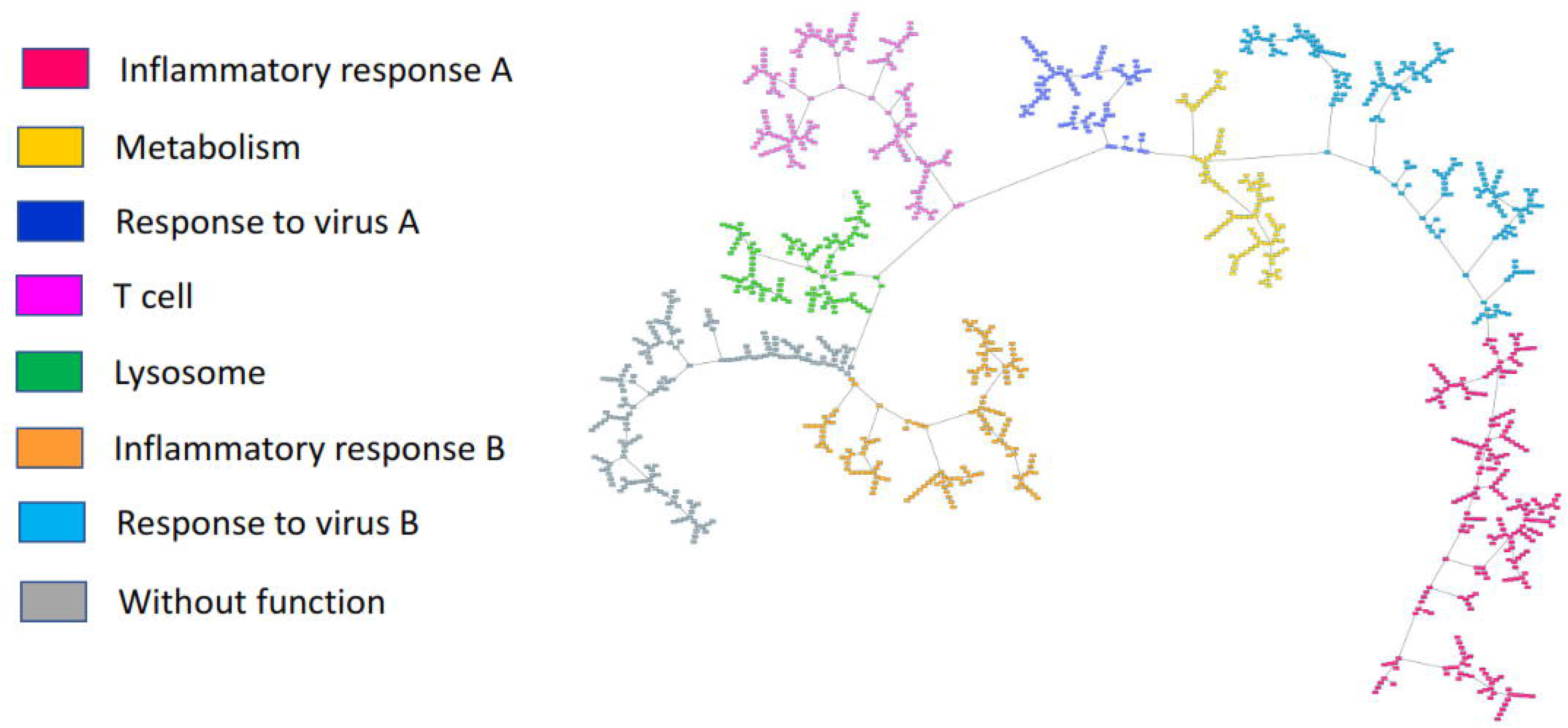
Probabilistic graphical model based on the expression of 1,234 differential genes between healthy individuals and patients.

Inflammatory response A, lysosome ad metabolism functional nodes activities were significantly differential between healthy controls and patients. Strikingly, one of the nodes of inflammatory response had a higher functional node activity in healthy controls than in patients, and the other node of inflammatory response had a functional node activity higher in patients than healthy controls. The same tendencies were shown in response to virus nodes. Lysosome and metabolism had a higher functional node activity in patients than in controls. Finally, T cell functional node activity was higher in healthy individuals than in patients (Fig 2).

**Fig 2:**
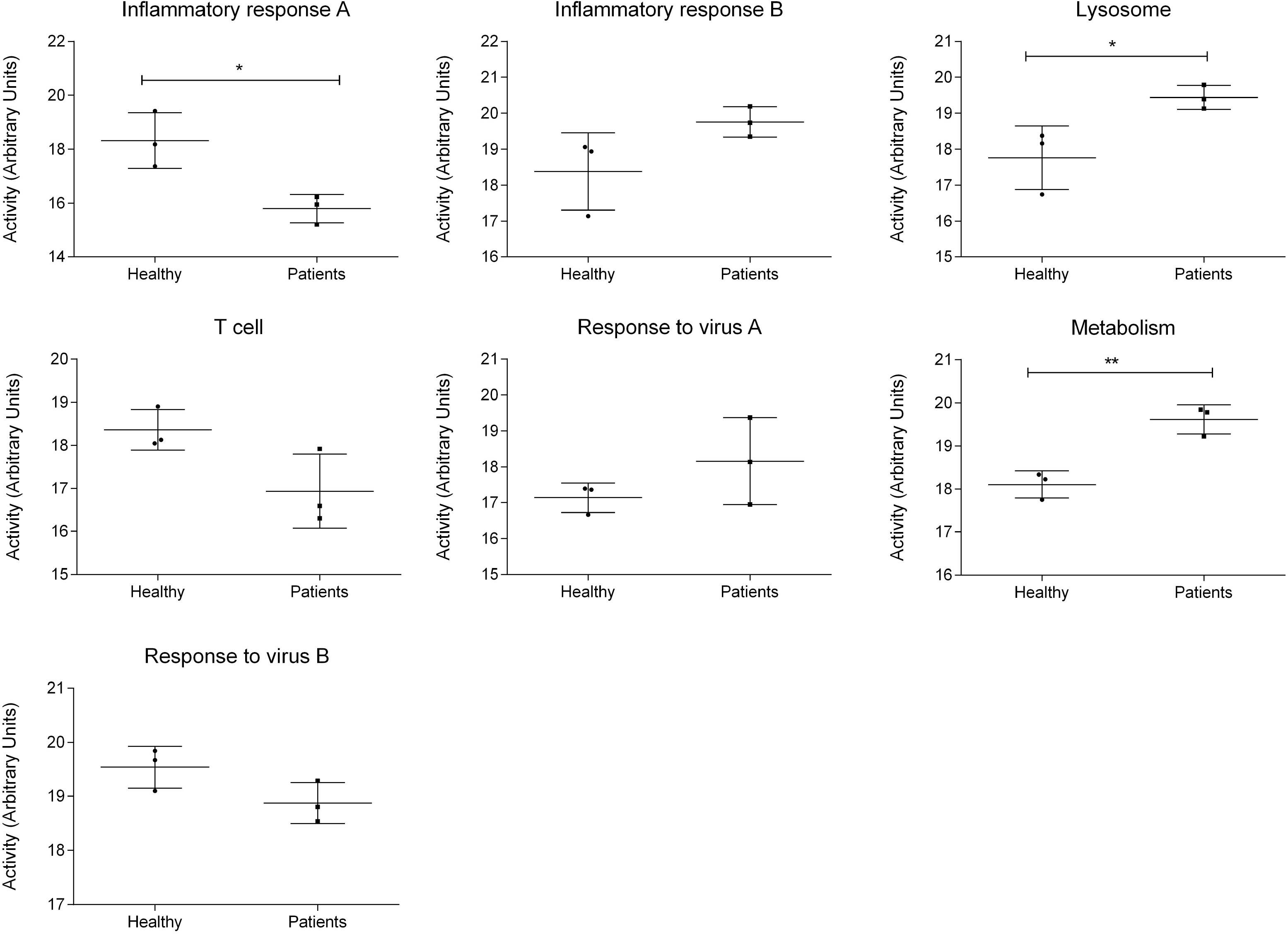
Functional node activities from the network based on the expression of the 1,234 genes defined as significantly differential by CuffDiff. In the Y axis the activity of the functional node in arbitrary units, understanding as the mean expression of those genes in each node that were related to the overrepresented function in the node. In the x axis, healthy controls and Covid-19 patients. **, ≤ 0.01; * ≤ 0.05.

#### Metabolism functional node

Metabolism node was composed of 102 genes, 19 of them related with metabolism pathways. This node contained the genes PKM (pyruvate kinase) and PDHB (pyruvate dehydrogenase), both implicated on glycolysis, and several ATPases from the mitochondrial complex, responsible for H+ transporting. There were also genes related to drug metabolism such as PAPSS1 or CES1.

#### Inflammatory response A functional node

This node included 236 genes, of which 18 were implicated in an inflammatory response. This node mostly comprised chemokines such as CXCR2, CCR3, CXCR1, CXCL1 or CXCL8, widely associated with SARS-CoV-2 infection.

#### Inflammatory response B functional node

This node included 129 genes, of which 17 were involved in inflammatory response. This functional node included three toll-like receptors, TLR2, TLR5 and TLR4. Among other functions, TLR2 promotes apoptosis in response to bacterial lipoproteins. TLR5 protein recognizes bacterial flagellin, the principal component of bacterial flagella. Additionally, TLR4 has been implicated in signal transduction events induced by lipopolysaccharide found in most gran negative bacteria. This node also contained AOAH and CD14, both genes implicated in response to bacterial lipopolysaccharides as well. Finally, this functional node included VNN1 which plays a suppressive role in influenza virus replication in human alveolar epithelial cells [7]. Therefore, this node is mostly related to the response to bacterial infections.

#### Lysosome functional node

Lysosome node included 112 genes, of which 12 were related to lysosomal processes. Most of these genes are lysosomal enzymes such as CTSH, NAGA or PLA2G15, but this node also included HPSE, which is the gene that encodes an enzyme that cleaves heparan sulfate proteoglycans to allow cell movement through remodeling of the extracellular matrix, or DRAM1, which encodes a lysosomal membrane protein that is requires for the induction of autophagy.

### Metabolic modeling

FBA was performed to characterize in depth metabolic alterations caused by SARS-CoV-2 infection (S2 File). Glutamate metabolism, methionine and cysteine metabolism, and tetrahydrobiopterin metabolism flux activities were differential between healthy controls and Covid-19 patients (Fig 3). In addition, several tendencies in other metabolic pathways such as TCA cycle and steroid metabolism that need to be confirmed were shown (S1 Figure).

**Fig 3:**
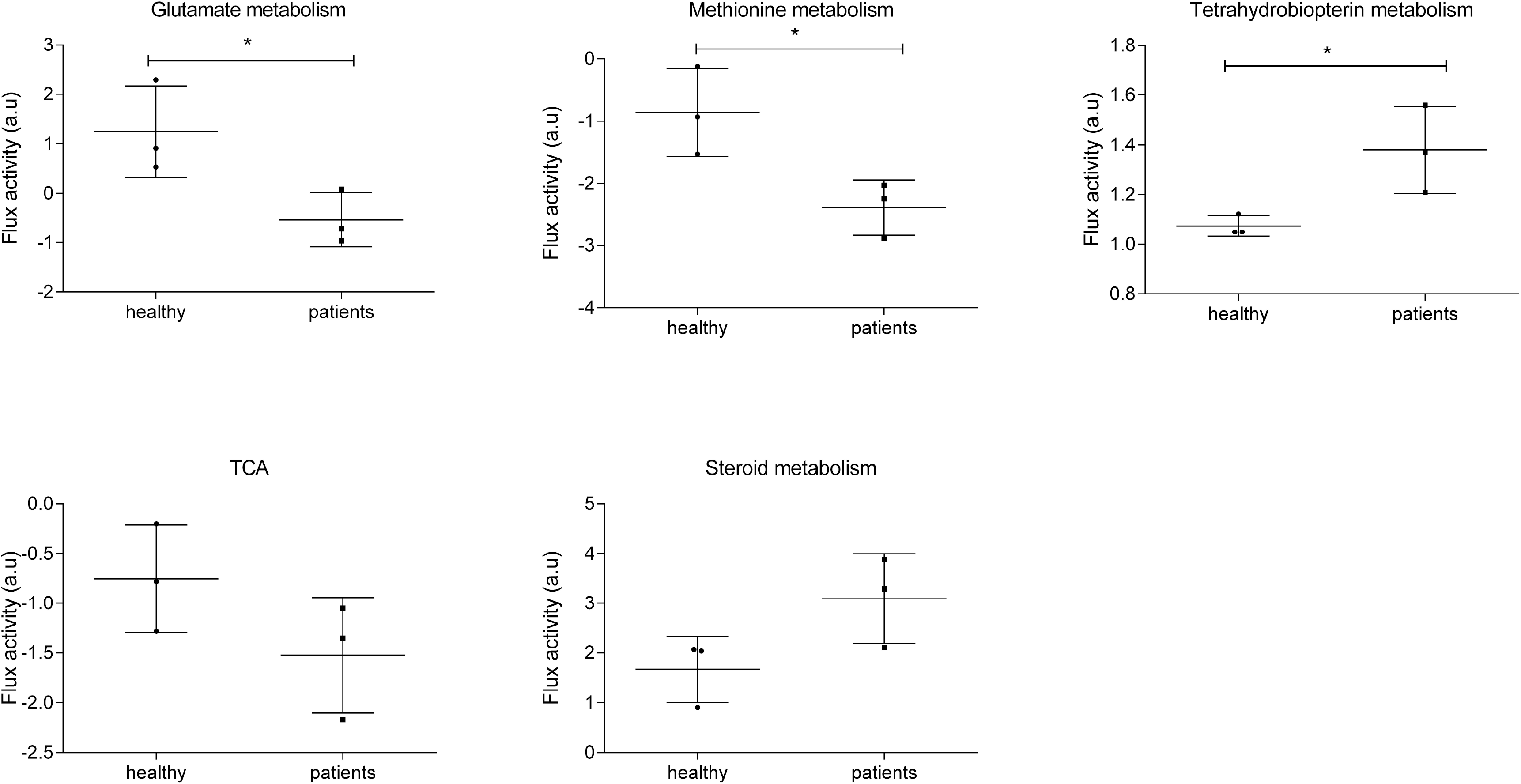
Differential flux activities between healthy controls and patients. a.u. = arbitrary units. * p < 0.05.

### Functional characterization based on the 2,000 most variable genes

We obtained an alternative PGM network, now based on the 2,000 most variable genes according to their SD and functionally characterized. The resulting network was divided into nine functional nodes: apoptosis, oxygen binding, blood coagulation, response to the virus, T cell, cell division, and three nodes related to an inflammatory response (Fig 4, S3 File).

**Fig 4:**
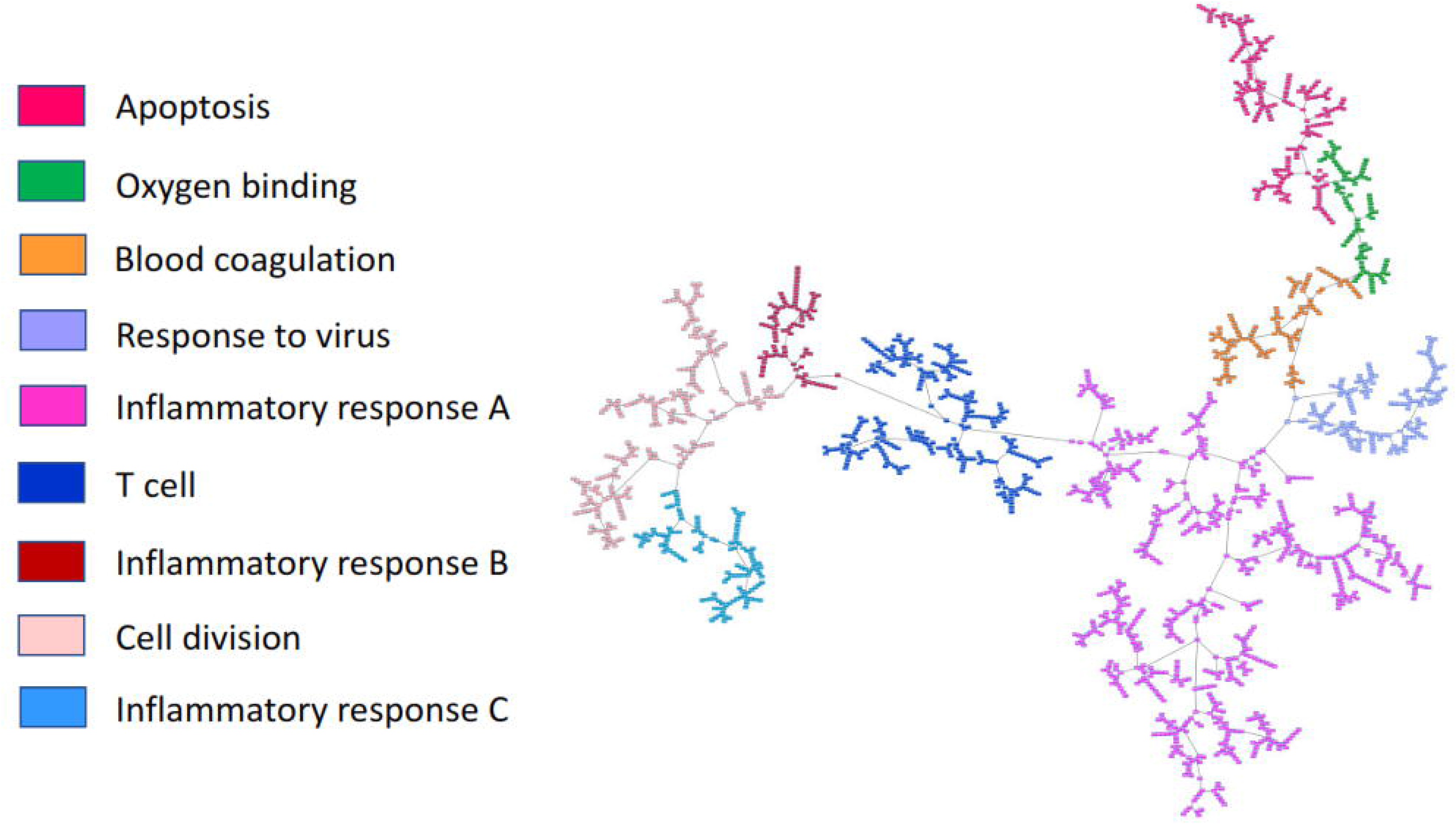
Probabilistic graphical model based on the expression of the 2,000 most variable genes.

Cell division and inflammatory response B functional node activities were differential between controls and patients. On the one hand, patients had a higher activity of response to virus, cell division, and one of the inflammatory functional nodes. On the other hand, healthy individuals had a higher activity of T cell and two out of the three functional nodes related to inflammatory processes (Fig 5).

**Fig 5:**
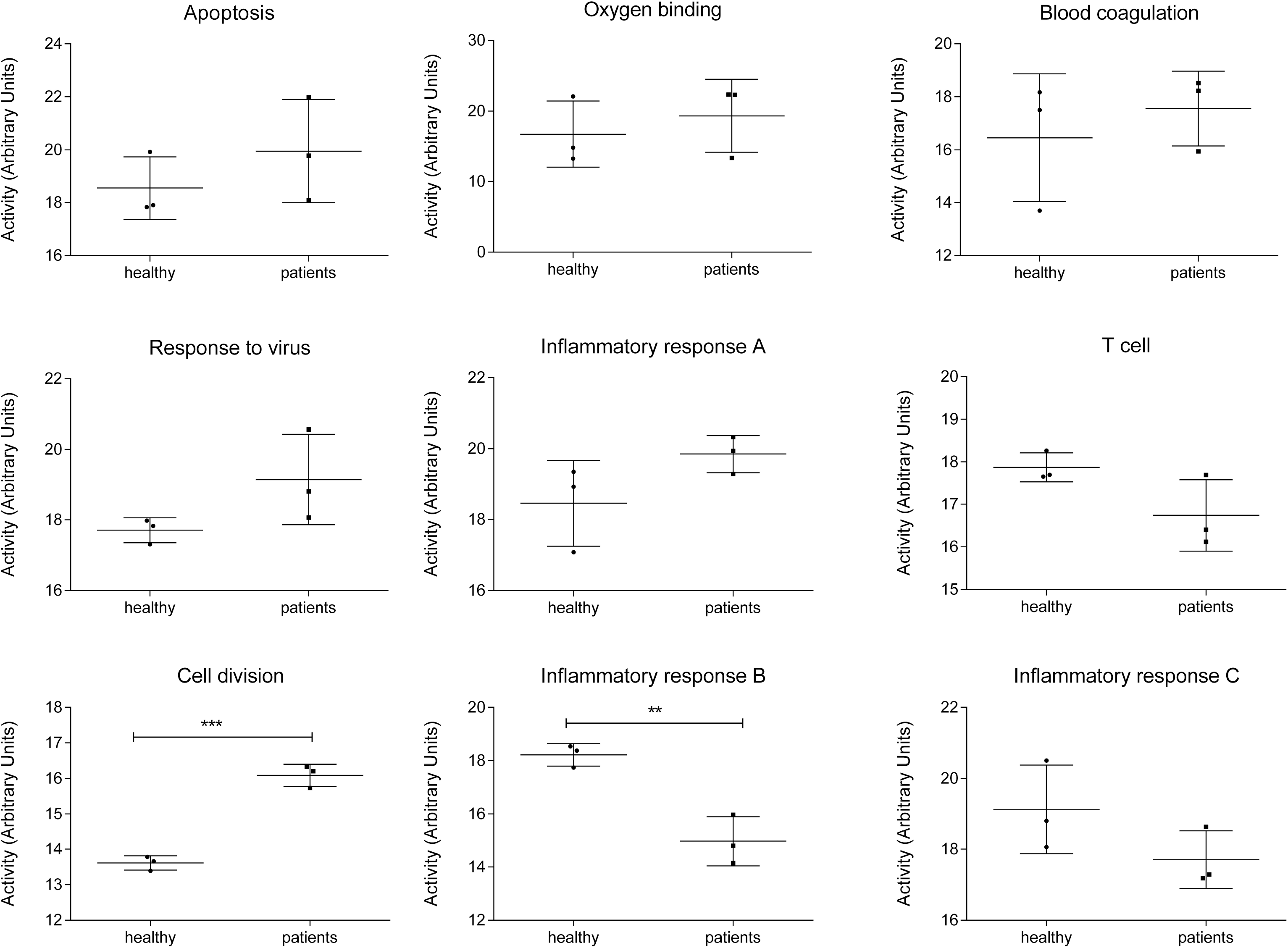
Functional node activities from the network based on the expression of the 2,000 most variable genes. In the Y axis the activity of the functional node in arbitrary units, understanding as the mean expression of those genes in each node that were related to the overrepresented function. In the X axis, healthy controls and Covid-19 patients. ***, ≤ 0.001; **, ≤ 0.01; * ≤ 0.05.

#### Inflammatory response A functional node

This node is composed of 77 genes, of which 28 were related to an inflammatory response. Most of these genes were cytokines, chemokines and toll-like receptors, whose function is the modulation of the inflammatory response. This functional node included toll-like receptors TLR6, TLR8, TLR5, TLR1, and TLR4.

#### Inflammatory response B functional node

This node was formed by 69 genes, 6 of them related to the inflammatory response. These six genes were CCR3, CCL4L2, TNFRSF18, NCR3, CCL5, and MS4A2. CCL5 and CCL4L2 are chemokine ligands, and CCR3 and MS4A2 are implicated in an allergic response.

#### Inflammatory response C functional node

This functional node had 77 genes, 6 of them related to inflammatory processes. These genes were CCL4, CXCR2, FPR2, IL1RAP, CXCL8, and ORM1, mostly of them chemokines.

#### T cell functional node

This functional node was composed of 210 genes, 6 of them related to T cells, more concretely with T cell receptors, including GATA3 gene, which plays a vital role in nasopharyngeal virus detection.

#### Cell division functional node

This node was composed of 141 genes, 5 of them involved in cell division. These genes were CDK1, CENPW, CCNB1, UBE2C, and CCNB2, mostly related to M-phase promoting factor complex and microtubules.

#### Response to virus functional node

This node had 130 genes, nine related to response to virus ontology. This node contained two genes whose proteins are induced by interferon, IFI44L and IFITM3.

## Discussion

SARS-CoV-2 infection has reached the category of a pandemic. Tremendous efforts have been made to find a suitable vaccine and to determine effective treatments but till the date there are neither of them [8].

We have re-analyzed the work of Xiong et al. [4] with a different functional inference approach. Coincidences between both results were expected. Xiong et al. described an up-regulation of genes related to cell cycle and cytokines in SARS-CoV-2 patients, which agreed with the higher functional node activity that we observed in cell division node and one of the inflammatory nodes, mostly composed by cytokines and chemokines. They also described a reduction of immune cells in blood patient samples, which may be related to the lower activity of T cell node in patients than in healthy controls.

Additionally, our analysis offered complementary information. For instance, in the network that characterizes differences between healthy individuals and SARS-CoV-2 patients based on the 1,569 differential genes identified by CuffDiff, two functional nodes related to inflammatory response were identified. Strikingly, inflammatory response A functional node activity was higher in patients than in healthy controls. This node was composed by cytokines and chemokines. However, inflammatory response B node activity, that is related to response to bacterial infections, was higher in healthy controls than in patients. SARS-CoV-2 coexist with a bacterial co-infection of *Mycoplasma pneumoniae* so the study of those genes related to the presence of a bacterial infection in these patients may be relevant [9]. Metabolism node showed also a higher functional node activity in Covid-19 patients than in healthy controls. The increase in glycolysis reactions implies an increase in Krebs cycle reactions as well and therefore in ATP production, essential for the virus replication [10, 11]. The differences on the metabolism functional node suggested that a deeper analysis of metabolism, as Flux Balance Analysis, could supply more detail information. Glutamate metabolism showed differences between controls and Covid-19 patients. Interestingly, an alteration in glutamate metabolism caused by another RNA virus, the HIV-1, has been previously described [11]. Moreover, it has been previously suggested that methionine plays a relevant role in viral replication of other coronaviruses [12]. No alterations in tetrahydrobiopterin metabolism have been previously described related to SARS-CoV-2 infection. However, it is remarkable that tetrahydrobiopterin is a NO synthase cofactor which is involved in immune regulation and inflammation processes. It has been described that a blockade of tetrahydrobiopterin synthesis annuls T-cell mediated autoimmunity and allergic inflammation. On contrast, higher levels of tetrahydrobiopterin increase CD4 and CD8 responses [13]. It has also been described that acute inflammatory stimulation increases levels of plasma BH4, in parallel with increased IL-6 [14], which it has been widely associated with SARS-CoV-2 infection and severity [15, 16]. Recent articles where plasma samples from Covid-19 patients were analyzed by metabolomics have shown differences in metabolism caused by SARS-CoV-2 infection, especially in steroid, aminoacid and mitochondrial metabolism [17].

Lysosomes have been previously associated with coronaviruses. In 1984, a study described virus-containing electron-dense bodies in lysosomes of coronavirus-infected cells as a defense mechanism [18]. Moreover, a study done in murine hepatitis virus, a prototype to study coronaviruses, established that the virus depends on the lysosomal traffic for a proteolytic cleavage site in the S protein, necessary for the intracellular fusion and entry [19]. In addition, this node contains the HPSE genes which it has been previously associated with viral infection and its activation is associated with a production of pro-inflammatory factors [20].

On the other hand, in the network obtained for the 2,000 most variable genes, two functional nodes related to inflammatory response (inflammatory response A and inflammatory response B) were also identified. Inflammatory response A functional node was again mostly composed by cytokines and chemokines. Inflammatory response B functional node had a lower activity in SARS-CoV-2 patients. Interestingly, the inflammatory response B node was composed of genes related to allergic response and regulation of immunological self-tolerance. This fact may be related to the severe acute respiratory syndrome, associated with a dysregulation of the immune response [2].

In this inflammatory response B node is included CCR3, a chemokine highly expressed in eosinophils and basophils, and is also detected in TH1 and TH2 cells, as well in airway epithelial cells [21, 22]. This receptor may contribute to the accumulation and activation of inflammatory cells in allergic airway and it is also known to be an entry co-receptor for HIV-1. MS4A2 is also implicated in allergic processes [23]. Therefore, this node seems to be more related to the self-control of the inflammatory response instead of the other inflammatory functional nodes, more related to chemokines and cytokines.

Cell division functional node had a significantly higher activity in Covid-19 patients than in controls. This node is mainly composed by genes related to M-phase and mitosis process, which may be related to viral infection. An accumulation of G2/M phase cells in other coronaviruses has been previously described in order to promote favorable conditions for viral replication [24].

As expected, functional nodes related to response to the virus were relevant in both networks. In the case of the network built based on the 2,000 most variable genes, this functional node was mainly related to interferon response. Remarkably, this node included IFITM3 gene, which codifying sequence is associated with immunity to other well-known viruses such as influenza A or dengue virus [25, 26]. IFITM3 protein has been described as related to the entry of MERS-CoV and SARS-CoV[27]. The first response to a viral infection of the immune system is mediated by interferons so it seems logical that these genes were overexpressed in patients infected by SARS-CoV-2. Additionally, interferon-mediated response has been associated with severe cases of Covid-19, so a study of the genes included in this functional node in a large cohort with different grades of severity of Covid-19 may be interesting [28].

In addition, in the T cell functional node appeared GATA3 gene which has been previously related to nasopharyngeal virus infections [29]. Since SARS-CoV-2 presents mainly respiratory tropism, GATA3 may play an essential role.

Our study had some limitations. Probably the most important one was that the reduced number of samples limited the statistical power and the information that could be obtained by functional analyses. A larger number of samples will be useful to deepen into the molecular characterization of this disease. Also, a study based on a larger cohort stratified according the severity of the disease could be of much interest as it may help define how functional modules vary in relation to the virulence of the infection.

In this study, some previously not described relevant processes in SARS-CoV-2 pathogenesis such as bacterial inflammatory response processes, tetrahydrobiopterin metabolism or allergic processes, were proposed. In the absence of treatments for these patients, molecular characterization of the disease could be helpful to improve the understanding of the mechanisms of the disease and to define targetable processes. The application of these type of analyses in larger cohorts may be useful not just to determine therapeutic targets but also to define predictors of immune response to infection. Therefore, these results may be relevant to propose new therapeutic treatments in the future.

## Materials and Methods

### Patient cohort

Three samples from peripheral blood mononuclear cells (PBMCs) from three patients infected with SARS-CoV-2 and three samples from healthy controls were analyzed. These samples are all from the work of Xiong et al. [4] and raw data can be downloaded from SRA database.

### Processing of RNA sequencing data

Before processing fragments per kilobase of exon model per million of reads (FPKM) data, we checked their quality using FastQC (v0.11.9, Brabaham, UK). Reads longer than 100 nt showed the presence of Illumina adapter sequences which were removed by trimming using Prinseq [30]so all samples were matched to 2×100 format. Then, reads were mapped against the human genome (GRCh38.96) using TopHat, using an estimated paired-end inner size of 25 and finally FPKM data were obtained using CuffDiff. All these programs were accessed using the integrated GPRO suite (Biotechvana, Valencia, Spain) [31].

After FPKM processing, Perseus v1.6.5 software was used to filter RNAseq data [32]. Log2 was calculated and only those genes with at least 50% of the detectable readings were used for the subsequent analyses.

### Probabilistic graphical models

2,000 most variable genes were selected according to their standard deviation (SD) of expression across the series and used to build a PGM network. RNAseq expression data was used without other a priori information.

The resulting network was divided into functional nodes by gene ontology analyses. These gene ontology analyses were performed in DAVID webtool v8 using “Homo sapiens” as background and KEGG, Biocarta and GOTERM-FAT as categories [33].

The same analysis pipeline was used to characterize the differential genes defined by CuffDiff, i.e. a network was built using the genes defined as significantly differential between healthy controls and patients.

These analyses was done using *grapHD* package [34] and R v3.2.5. Network visualization was done in Cytoscape [35]. PGMs were built in two steps, first, the spanning tree with the maximum likelihood was found, and then, the edges were refined based on the minimization of the Bayesian Information Criterion (BIC) [36].

### Statistical analysis

Functional node activities were calculated as previously described [6]. Briefly, the mean expression of those genes of each node related to the overrepresented function in this node was calculated. Then, functional node activities were compared between healthy individuals and patients using a T-test.

### Flux Balance Analysis and metabolic models

Flux Balance Analysis (FBA) allows metabolic modeling from gene expression data. It is widely used in microbiology and cancer [37]. The complete human metabolic reconstruction Recon 3D was used to perform these analyses. It contains 10,600 reactions, 5,835 metabolites and 5,939 Gene-Protein-Reaction rules (GPRs), which contain information in the form of Boolean expressions about which genes are involved in each metabolic reaction. GPRs were solved using a modification of Barker et al. algorithm [38, 39], solving “AND” expressions as the minimum and “OR” expressions as the sum. Then, the obtained values were introduced as the reaction bounds by a modified E-flux algorithm based on the Max-min function [39, 40]. Finally, FBA was solved using COBRA Toolbox library v2.0 [41] and MATLAB.

The 10,600 metabolic reactions are grouped into 103 metabolic pathways or subsystems. In order to compare metabolic activity between controls and Covid-19 patients, flux activities were calculated as previously described as the sum of fluxes of the reactions contained in a concrete metabolic pathway [5, 42]. To compare flux activities between control and patients a T-test was used.

## Supporting information

S1 File: Genes included in the probabilistic graphical model based on the expression of 1,234 differential genes between healthy individuals and patients.

S2 File: Genes included the probabilistic graphical model based on the expression of the 2,000 most variable genes.

S3 File: Flux Balance Analysis results.

